## Supplemental Information for "Reference compounds for characterizing cellular injury in high-content cellular morphology assays"

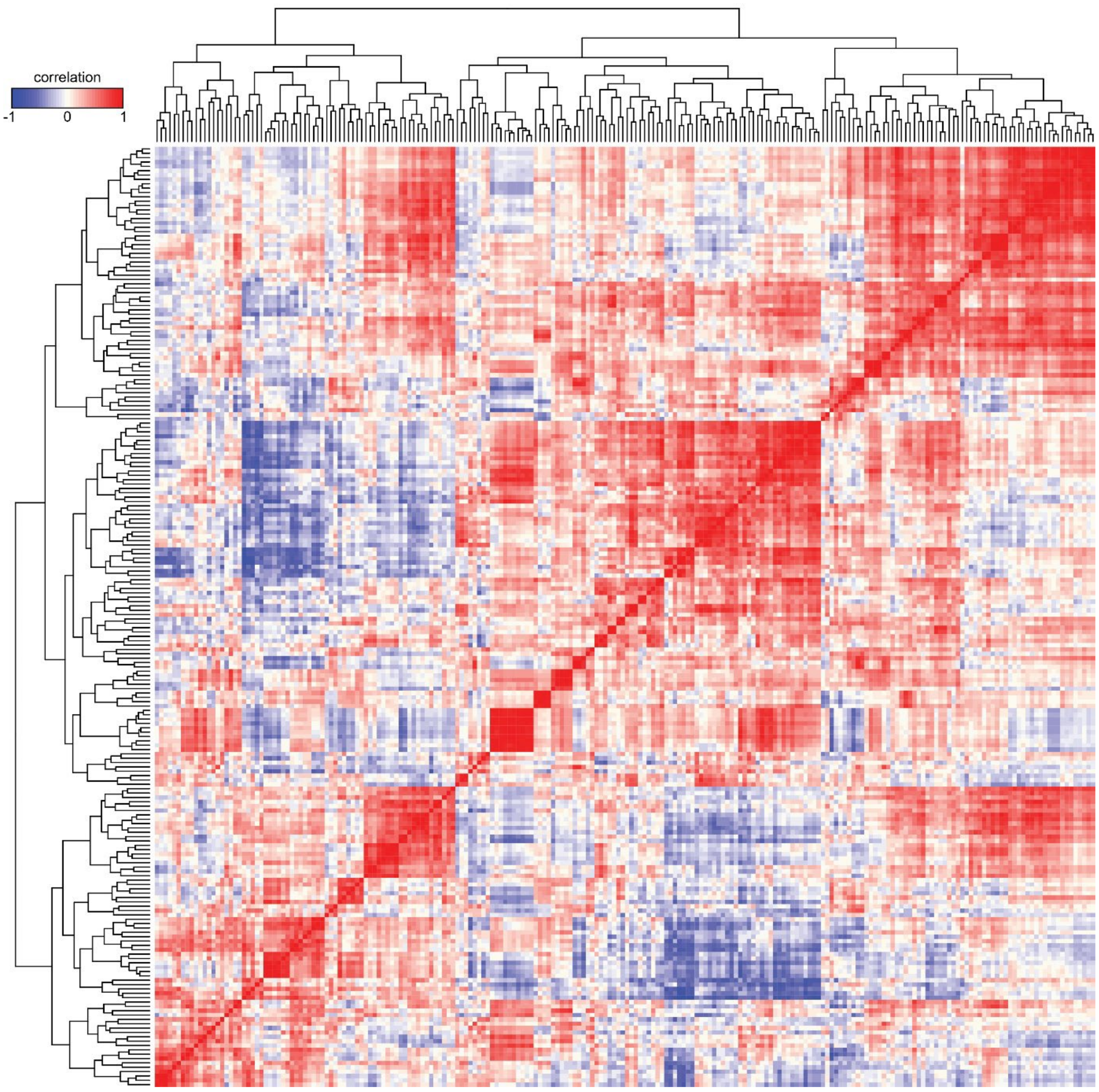


**Supplementary Figure 1. Summary of cell painting compound clustering.** Compounds (**1-171, 455-501**) were sorted by unsupervised hierarchical clustering. Data are from 24 h compound treatment times.

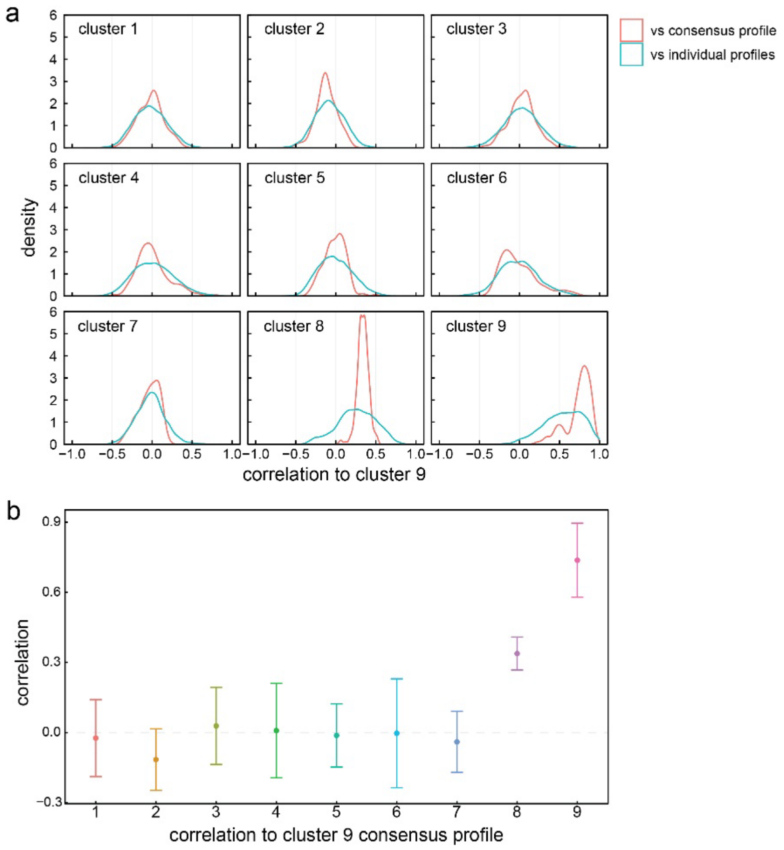


**Supplementary Figure 2. Calculation of cellular injury phenotype.** (a) The cell injury phenotype is calculated as the average signature of cluster 9, where the average value for each feature across all cluster 9 members is taken, and the vector comprising these average values is used as the cell injury phenotype. This average profile can be considered the central point of the cloud of cell injury phenotypes in multidimensional space. While the profiles of compounds belonging to cluster 9 were somewhat diverse, their correlation to this consensus cell injury phenotype was more robust compared to the pairwise correlation to all compounds belonging to cluster. There is a significant improvement in the consistency of these correlations when comparing the consensus cell injury phenotype to individual compounds belonging to cluster 8. (b) Comparing the correlations of all compounds to this consensus cell injury phenotype, we find that those belonging to cluster 9 have the highest correlations to the consensus profile, suggesting that this metric can be used to predict MLI compounds that might also induce similar cell injury phenotypes. Data are mean ± SD.

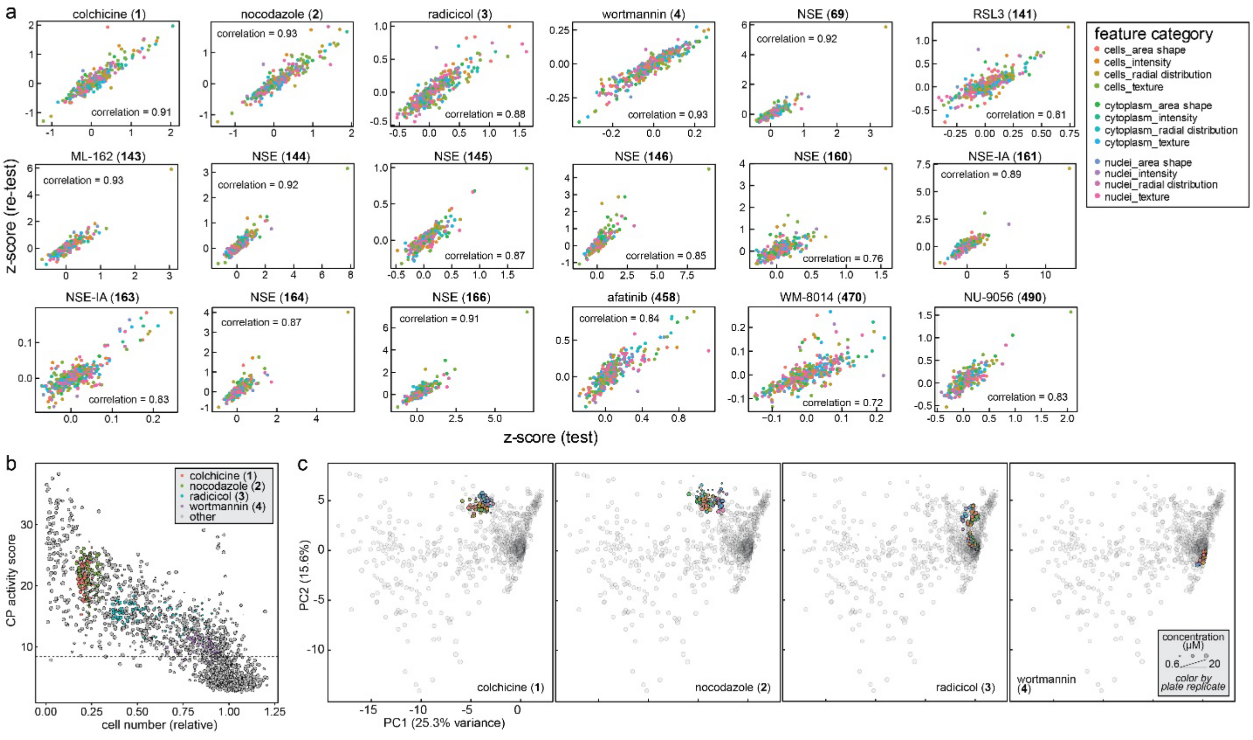


**Supplementary Figure 3. Cell painting phenotypes show acceptable reproducibility.** (a) Select compounds show high correlations across independent CP experiments (test and re-test). The scatterplots depict the z-scores of each cellular feature, with points colored according to the feature category. (b, c) CP positive controls **1-4** show reproducible activities, cell numbers, and PCA clustering after 24 h treatment times across 13 compound plates (pooled from 2 independent experiments, 4 replicate plates per compound plate).

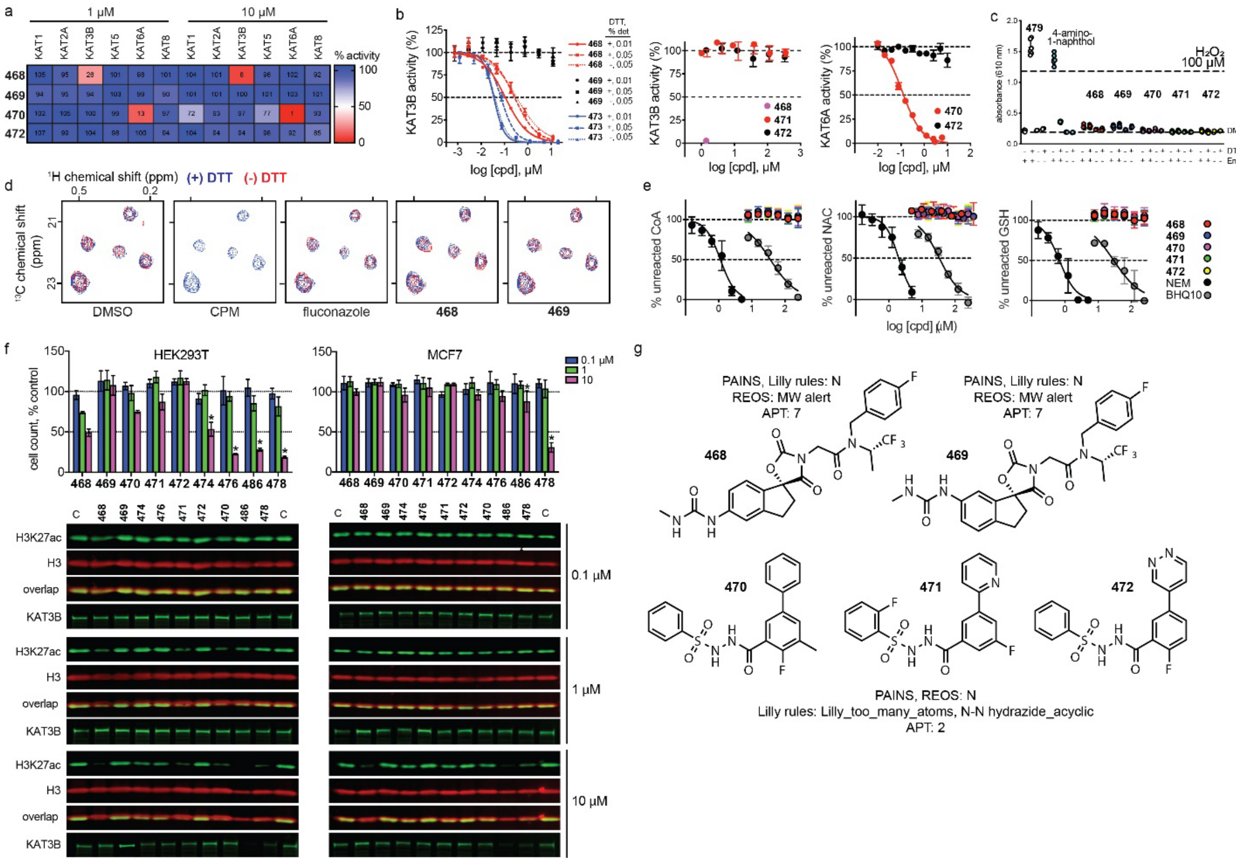

**Supplementary Figure 4. Next-generation KAT inhibitors are high-quality chemical probes with minimal common assay interferences.** (a, b) Compounds **468** and **470** are potent and selective cell-free inhibitors of KAT3B and KAT6A, respectively. (c) Compounds **468**-**472** do not produce detectable levels of hydrogen peroxide. (d) Compounds **468** and **469** are ALARM NMR-negative, suggesting they are not grossly reactive with protein thiols. (e) Compounds **468-472** do not react with the non-proteinaceous thiols CoA, GSH, or NAC by fluorometric thiol reactivity assay. (f) Top: prototypical interference compounds, but not next-generation KAT inhibitors, disrupt HEK293T and MCF7 cell viability. Bottom: the next-generation KAT inhibitor **468** and some prototypical interference compounds (**478**, **486**) decrease H3K27ac levels. (g) Compounds **468-472** exhibit do not exhibit red flags by cheminformatics analysis for substructure PAINS, REOS, Lilly alerts (FAF-Drugs4, accessed 11 July 2018) and Abbott Physicochemical Tiering (Pipeline Pilot, BIOVIA, version 9.0.2.1).

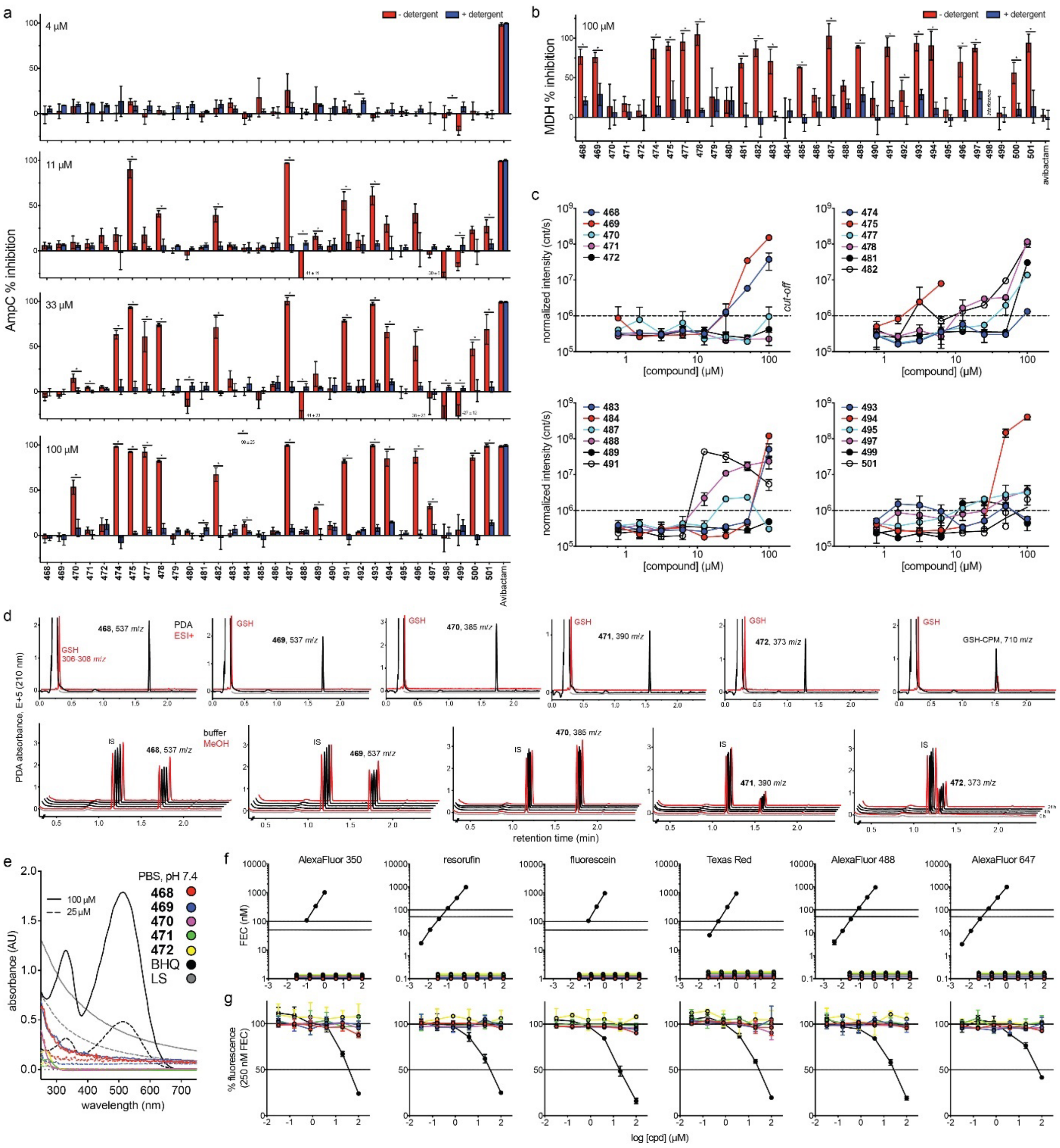

**Supplementary Figure 5. Next-generation KAT inhibitors show minimal cell-free interferences by additional counter-screens.** (a, b) Many historical KAT inhibitors show detergent-sensitive inhibition of AmpC β-lactamase and MDH. Detergent-sensitive differences in inhibition are a hallmark of compounds that form colloidal aggregates. (c) Several KAT inhibitors form detectable colloidal aggregates by DLS. (d) Compounds **468**-**472** do not form grossly detectable adducts with GSH (top) and are grossly stable in assay-like conditions by UPLC-MS (bottom). (e) Compounds **468**-**472** do not significantly absorb visible light in PBS. PC, BHQ-10 carboxylic acid ($\lambda_{max}$ = 516 nm); LS, creamer light scattering control (1 and 0.5 mg mL$^{-1}$). (f, g) Compounds **468-472** do not fluoresce or quench fluorescence in six common fluorophore channels. Dotted lines, U/LLOQ. PC auto-fluorescence, independent fluorophore sample; PC quenching, BHQ-10.

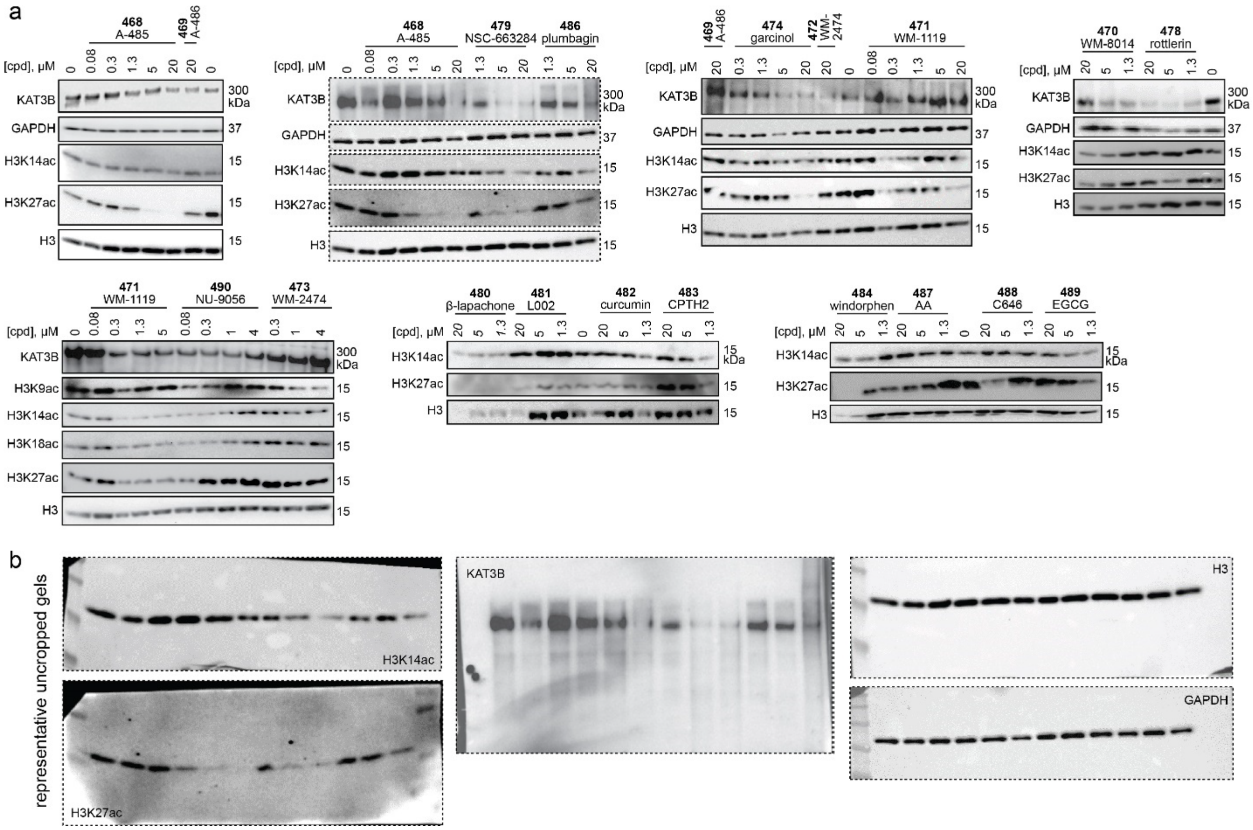


**Supplementary Figure 6. Next-generation KAT inhibitors and prototypical nuisance compounds produce overlapping but distinct cellular histone acetylation phenotypes in cell painting-like conditions.** (a) U-2 OS cells were treated with compounds for 24 h at the indicated concentrations under experimental conditions mimicking the CP assay. Compound **468** (but not inactive analog **469**) potently reduced cellular H3K27ac levels (approximate $EC_{50}$ = 1 μM) and did not modulate H3K14ac or KAT3B levels. The KAT6-specific inhibitors **470** and **471** mildly decreased cellular H3K9ac, H3K14ac, and H3K27ac levels. Note that these compound predominately target KAT6A, which acetylates primarily H3K23. Consistent with other profiling trends (**Supplementary Figure 3**), several prototypical interference compounds and historical KAT inhibitors decreased H3K27ac levels, while also markedly reducing KAT3B levels in U-2 OS cells. *Italicized concentrations*, notable cell death. Histone H3 and GAPDH, H3Kac and KAT3B normalization controls, respectively. (b) Represenative uncropped gel images.

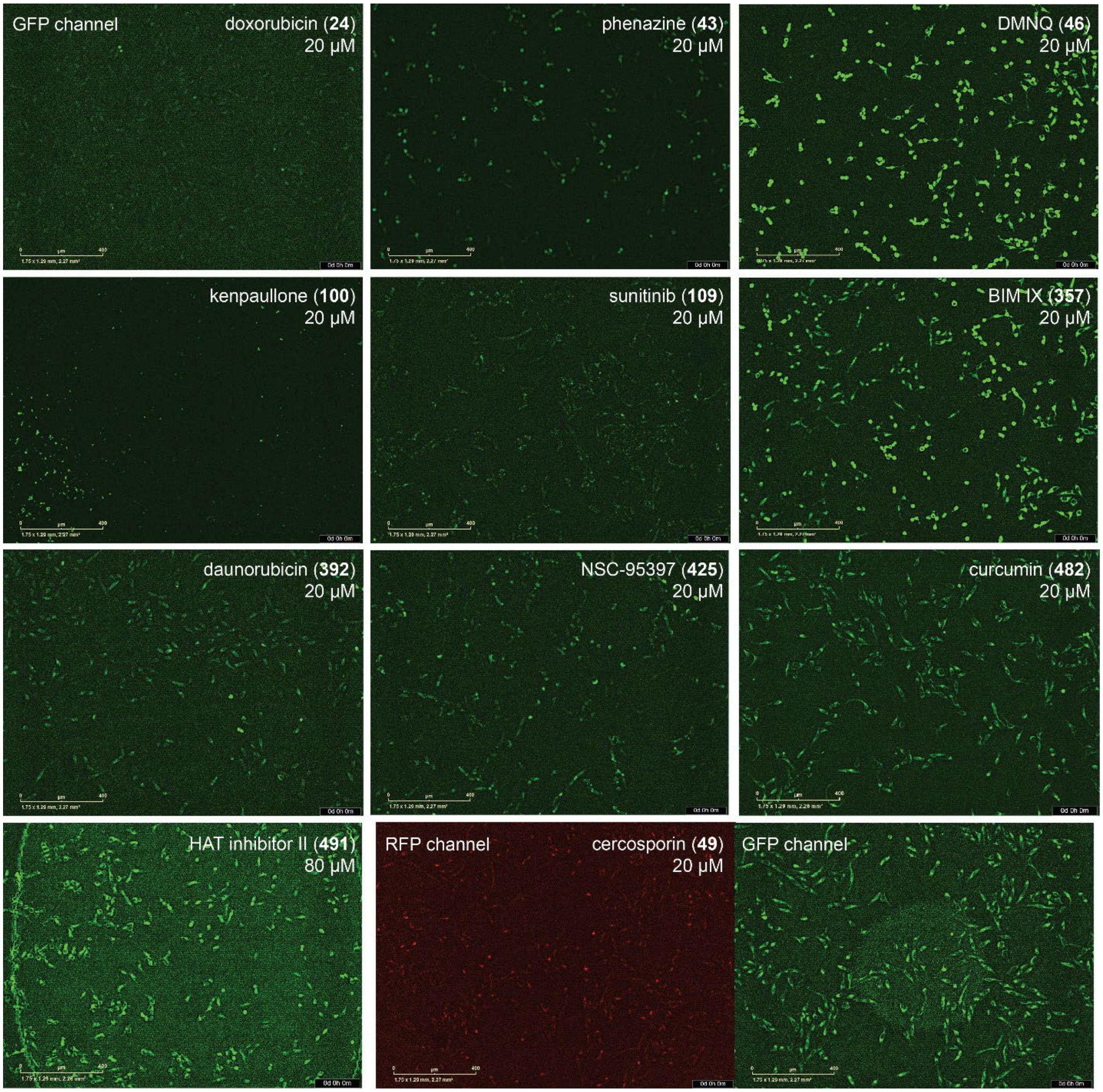


**Supplementary Figure 7. Compound-dependent artifacts in live-cell imaging identified by reagent-free counter-screen.** Representative examples of auto-fluorescent compound artifacts in live-cell imaging. U-2 OS cells were treated with compounds for 1 h in the absence of cellular health reagents, followed by imaging and processing otherwise identical to the primary live-cell cellular health assay. Note that most fluorescent compound artifacts interfered with the GFP (green) rather than the RFP (red) channel. Image scales: 400 μm.

**Supplementary Table 1. Sources of KAT inhibitors.**

| Compound | Name | Source | Notes |
|---|---|---|---|
| **468** | A-485 | Abbvie | *Via* SGC |
| **469** | A-486 | Abbvie | *Via* SGC |
| **470** | WM-8014 | Monash | Synthesized in-house using published procedures[1,2]. |
| **471** | WM-1119 | Monash | |
| **472** | WM-2474 | Monash | |
| **473** | Lys-CoA | NCATS | |
| **474** | Garcinol | Cayman Chemical | Cat # 10566 |
| **475** | Isogarcinol | Cayman Chemical | Cat # 21164 |
| **476** | Dahlin-JMC2015-6a | UMN | |
| **477** | Embellin | Cayman Chemical | Cat # 11838 |
| **478** | Rottlerin | Cayman Chemical | Cat # 12006 |
| **479** | NSC-663284 | Cayman Chemical | Cat # 13303 |
| **480** | Beta-lapachone | Santa Cruz Biotechnology | Cat # sc-2000875 |
| **481** | L002 | Cayman Chemical | Cat # 17778 |
| **482** | Curcumin | Sigma-Aldrich | Cat # 08511 |
| **483** | CPTH2 | Cayman Chemical | Cat # 12086 |
| **484** | Windorphen | Sigma-Aldrich | Cat # SML0899 |
| **485** | Windorphen-NC | Calbiochem | Cat # 509164 |
| **486** | Plumbagin | Cayman Chemical | Cat # 14314 |
| **487** | Anacardic acid | Cayman Chemical | Cat # 13144 |
| **488** | C646 | Cayman Chemical | Cat # 10549 |
| **489** | EGCG | Cayman Chemical | Cat # 70935 |
| **490** | NU-9056 | Tocris | Cat # 4903 |
| **491** | HAT Inhibitor II | Cayman Chemical | Cat # 19835 |
| **492** | Rtt109 Inhibitor | Enamine Ltd. | Cat # Z14250979 |
| **493** | MG149 | Cayman Chemical | Cat # 22135 |
| **494** | EML425 | EMD Millipore | Cat # 534354 |
| **495** | MB-3 | Sigma-Aldrich | Cat # M2449 |
| **496** | SPV-106 | Sigma-Aldrich | Cat # SML0154 |
| **497** | Gossypol | Cayman Chemical | Cat # 14482 |
| **498** | CTK7A | Sigma-Aldrich | Cat # 382115 |
| **499** | TH 1834 | Axon Medchem | Cat # 2339 |
| **500** | Nordihydroguaiaretic acid | Sigma-Aldrich | Cat # 74540 |
| **501** | CAY10669 | Cayman Chemical | Cat # 10974 |

**Supplementary Note 1. Next-generation KAT inhibitors favorable by cheminformatics analyses.** Literature analyses and scrutiny of the chemical structures of reported cell-active KAT3B and KAT6 tool compounds **468-472** raised minimal alerts. We note that substructure filters and calculated properties are only one of many tools for compound prioritization. None of these compounds were flagged as PAINS, though **468** and **469** were flagged by REOS filters for their higher MW. Lilly substructure filters flagged **470-472** for their hydrazide moieties (**Supplementary Figure 3**)[3-5]. Acylsulfonylhydrazides (acylsulfonohydrazides) have been viewed as problematic because of (1) the metabolic liability of the *N-N* bond, which can undergo facile cleavage or oxidation, (2) evidence that they form metal complexes, and (3) they can undergo facile nucleophilic attack under metal catalysis[6,7]. Compounds **470-472** showed favorable physicochemical properties by Abbott Physicochemical Tiering (APT), while **468** and **469** were tiered as less favorable because they are not Rule-of-Five compliant (**Supplementary Figure 3**)[8]. The latter is less concerning, as **468** has already been shown to be orally bioavailable[9]. Literature analysis showed that the oxazolidinedione core of **468** and **469**, although one might speculate might be mildly electrophilic with the potential to therefore be promiscuous, was not a frequent moiety reported in screening publications. Indeed, it is present in potent, selective, and convincing mineralcorticoid receptor antagonists, as well as the advanced kinase inhibitor, pegcantratinib[10,11]. Similarly, the sulfonoaroylhydrazide of **470-472**, while it could be speculated might bear a relatively electrophilic carbonyl group, did not appear significantly in screening literature and indeed has been reported by Pfizer to furnish potent inhibitors of human branched-chain amino transferase, with reasonable pharmacokinetic properties[12].

**Supplementary Note 2. Compounds 468-472 show potent, on-target cell-free inhibition of their reported KAT targets** (refer to **Supplementary Figure 2b**). Compound **468** is a potent inhibitor of KAT3B activity ($IC_{50}$ = 109 nM; complete response; non-cooperative Hill slope of 0.89, 95% CI 0.72 to 1.06) while **469** showed minimal activity. This is consistent with original reports ($IC_{50}$ = 9.8 and 60 nM), which were obtained by TR-FRET and SPA assays, respectively[13]. Neither excess electrophilic scavenging agent dithiothreitol (DTT), nor nonionic detergent grossly altered the potency of **468**, suggesting it is not acting by thiol-reactivity or aggregation[14,15]. These potencies are several-fold weaker than Lys-CoA (**473**), a well-behaved yet cell-impermeable bi-substrate analog used as a positive control compound[2,16]. As expected for KAT6-specific inhibitors, neither **471** nor **472** showed appreciable activity versus KAT3B activity. Compound **470**, showed potent *in vitro* inhibition versus KAT6A in the SPA format ($IC_{50}$ value of 114 $\pm$ 11 nM; complete response; non-cooperative Hill slope of 1.0). This is consistent with the original report ($IC_{50}$ = 8 nM) obtained by an AlphaScreen assay.

Differences in cell-free $IC_{50}$ values between the original report and this study are likely due to differences in assay design. For KAT3 inhibitors **468** and **469**, notable differences between the EMSA and previously published assay results include enzyme concentrations (0.2 versus 150 nM KAT3B, respectively), substrate concentrations (0.5 versus 1.0 $\mu$M acetyl-CoA, respectively), and protein construct (KAT3B BHC domain aa 1036-1822 versus KAT3B aa 1284-1673, respectively)[13]. For KAT6A/B inhibitors **470-472**, notable differences between the SPA and previously published assay results include protein construct (KAT6A aa 497-780 versus aa 472-793, respectively), substrate concentrations (15 versus 5 $\mu$M acetyl-CoA, respectively), reaction buffer components, assay temperature, and reaction times[1].

**Supplementary Note 3. Status of KAT3 in U-2 OS cells.** There are 3 and 4 copies of KAT3A and KAT3B, respectively, and no KAT3 mutations present in U-2 OS cells according to the Cancer Cell Line Encyclopedia[17]. The *KAT3B* gene is not under-expressed in osteosarcoma cells according to the Broad Institute DepMap[18]. The protein KAT3B is also expressed in U-2 OS cells according to the MaxQB proteomics database[19].

**Supplementary Note 4. Plate layout of typical cell painting experiment.** Each 384-well CP plate contains 116 solvent/negative control wells (DMSO), four positive control compounds (**1**-**4**; colchicine, nocodazole, radicicol, wortmannin), and 168 wells available for test compounds. Most compounds were profiled at six compound concentrations, diluted row-wise (e.g., C3 to C8, 20 to 0.625 μM final concentrations, respectively).

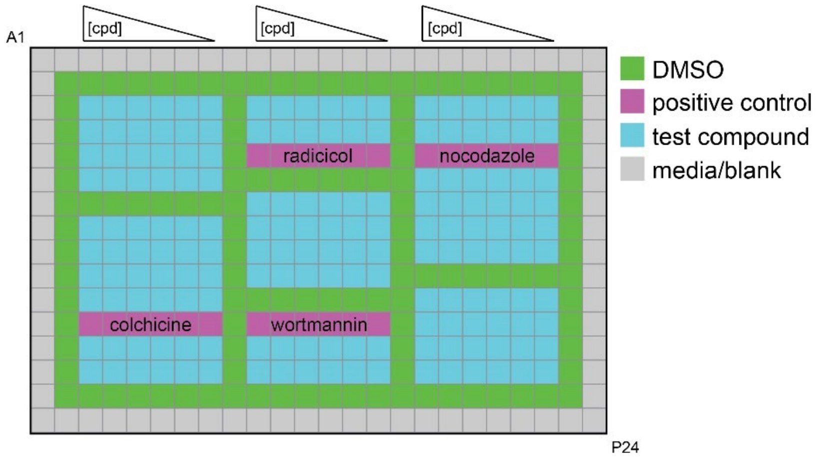
